# Impending Regeneration Failure of the Iucn Vulnerable Borneo Ironwood (*Eusideroxylon Zwageri*)

**DOI:** 10.1101/406025

**Authors:** Lan Qie, Alexander D. Elsy, Ashley Stumvoll, Magdalena Kwasnicka, Anna L. Peel, Joseph A. Sullivan, Maisie S. Ettinger, Alasdair J. Robertson, Jeanelle K. Brisbane, Amber L. Sawyer, Yan N. Lui, Siew Ngim Ow, Matteo Sebastianelli, Bartosz Majcher, Muying Duan, Hannah Vigus, Grace Pounsin, Reuben Nilus, Robert Ewers

## Abstract

The regeneration of many climax species in tropical forest critically depends on adequate seed dispersal and seedling establishment. Here we report the decreased abundance and increased spatial aggregation of younger trees of the Borneo ironwood (*Eusideroxylon zwageri*) in a protected forest in Sabah Malaysia. We observed a high level of seedling herbivory with strong density dependence, likely exacerbated by local aggregation and contributing to the progressively shrinking size-distribution. We also note the largely undocumented selective herbivory by sambar deer on E. *zwageri* seedlings. This study highlights the combined impact of altered megafauna community on a tree population through interlinked ecological processes and the need for targeted conservation intervention for this iconic tropical tree species.

## INTRODUCTION

The loss of megafauna due to climate change, habitat alteration and hunting has a strong impact on plants that are dependent on large-bodied animals for seed dispersal, and this effect is particularly strong in tropical forests (Corlett 2010, Galetti et *al.* 2017). Many tree species with large fruits have lost the animal dispersers they coevolved with, and consequently suffer reduced seed dispersal and increased spatial aggregation of seedlings, leading to lower survival and reduced gene flow (Harrison et *al.* 2013, Galetti *et al.* 2017). Such trees are typically late-succession to canopy species (Harrison *et al.* 2016), and their regeneration is crucial to the future of intact tropical forests and restoration efforts to accelerate succession in disturbed forests.

Borneo ironwood *(Eusideroxylon zwageri)* is a classic example of rainforest climax tree species facing regeneration challenges. *E. zwageri* is distributed in eastern and southern Sumatra, Bangka, Belitung, Borneo, the Sulu archipelago and Palawan. In Borneo, *E. zwageri* was formerly a common component of the main middle story in the mixed dipterocarp forest. It is a prized timber species, renowned for its extraordinary strength, durability and rot resistance, and it is a cultural keystone species to the indigenous people of the region (Franco *et al.* 2014). Despite being long-living (>1000 years), due to over-exploitation (Peluso 1992) and slow natural regeneration (typically requires >100 years to reach 30 cm diameter) (Irawan 2005), *E. zwageri* has become scarce across its distributional range and is classified as Vulnerable in the IUCN Red List (1998). The species produces large drupaceous fruits that measure 10-18 cm x 5-10 cm, and contain a single large seed, measuring 7-15 cm × 4-7 cm, with a very hard seed coat (Irawan 2005). Water dispersal for the heavy seeds of *E. zwageri* is possible but limited as the species often occur away from rivers (Irawan 2005). The large fruit with edible fleshy mesocarp and nut are strong indications that *E. zwageri* is dispersed by animals, although evidence of this is anecdotal and mainly refers to porcupines (Kostermans *et al.* 1994, Phillipps & Phillipps 2016). Only megafauna (>44 kg in body mass) would be capable of long-distance dispersal of these large seeds (Galetti *et al.* 2017). It has been speculated that *E. zwageri* seeds were dispersed by the Sumatran rhinoceros *(Dicerorhinus sumatrensis)* (Phillipps & Phillipps 2016), formerly occurring throughout Borneo but now virtually extinct in the wild in Sabah, and down to the last few individuals, if any, still surviving in Borneo (Hance 2015). Despite the value and threatened status of *E. zwageri,* we still have very limited understanding of its ecology and, in particular, its natural regeneration from seeds.

Post germination, the seedling stage represents another bottleneck for the regeneration of a tree species under survival pressures such as pathogens and seedling herbivory, which are typically density dependent processes (Wang & Smith 2002). There have been anecdotal reports that one of the common herbivores in Bornean forests, the sambar deer *(Rusa unicolor),* preferentially browsed on *E. zwageri* seedlings. This would potentially exacerbate the regeneration challenge already faced by *E. zwageri.* Once stems survive the dynamic stage and reach 10 cm diameter the degree of spatial aggregation is expected to be stable across size classes (Condit *et al.* 2000). Therefore in a long-living *E. zwageri* population (oldest trees may be > 1000 years), spatial aggregation in younger versus older trees should reflect long-term change in the strength of seed dispersal. We conducted a survey of *E. zwageri* across all life stages from seedlings to adult trees in an old growth Bornean lowland forest and tested three hypotheses: 1) reduced regeneration in this local population will be reflected in its tree size distribution curve with a flat gradient at small size classes, or a unimodal shape (Halpin & Lorimer 2017), 2) there is increased spatial aggregation in the younger sub-population compared to the older sub-population of adult trees (≥ 10 cm diameter), and 3) seedlings herbivory damage has density dependence.

## MATERIALS AND METHODS

We conducted this study in January 2018 at Maliau Basin Conservation Area (4.74° N, 116.97° E, 256 m a.s.l.), Sabah, Malaysia. The survey area was in old growth forest on the south bank of the Maliau River near the edge of the conservation area, in the vicinity of a large buffer zone consisting of selectively logged and secondary forests. A 4-ha square plot was set up, within which a grid of 10 by 10 m subplots was established for tree surveys. All *E. zwageri* trees with DBH ≥ 1 cm were mapped and recorded to diameter classes in 10 cm intervals.

There were abundant *E. zwageri* seedlings in the area from the January 2017 fruiting event. Many of these had noticeable herbivory damage with young leaves missing from the top, characteristic of that by mammalian herbivores, and often resulting in complete defoliation. We surveyed seedlings (all stages with DBH < 1 cm) in a 1.35 ha area in the centre of the 4-ha plot, using 5 by 5 m subplots within the 10 by 10 m grid. In each seedling subplot, we counted *E. zwageri* seedlings in two categories: “surviving” if the apical meristem was intact, and “fatally damaged” if the apical meristem was eaten, including dead *E. zwageri* seedlings with evidence of herbivory damage. Leafless and dead *E. zwageri* seedlings were identified by their characteristic straight and reddish bare stem, often with the stony seed coat still attached at root. For each subplot we also recorded ground vegetation cover as a proxy for interspecific competition, estimated into four classes, 0-25%, 25-50%, 50-75% and 75-100%.

We quantified spatial aggregation of *E. zwageri* adult trees (DBH ≥ 10 cm) using the relative neighborhood density Ω_x_, defined as the average density of neighbours in the annulus at distance *x* for each focal tree, standardized by mean population density (Condit *et al.* 2000). We calculated Ω_x_ for the 0-100m distance range in 10 m steps, and compared this metric for the older and younger sub-populations, categorized as those above or below the median tree size respectively.

For *E. zwageri* seedlings we used a generalized linear model (GLM) with a binomial error distribution and a logit link function to test the effects of conspecific seedling density and vegetation cover on the proportion of fatal herbivory damage. All analyses were conducted in the R statistical computing environment (R Core Team, 2016).

## RESULTS

We recorded 90 *E. zwageri* trees (DBH ≥ 1 cm) in the 4 ha plot, among which 80 were adult trees (DBH ≥ 10 cm). As predicted from the reduced regeneration hypothesis the adult tree size distribution had a unimodal shape peaking at the 70 cm class (median) and decreasing towards smaller classes (Fig. 1a). All adult trees were aggregated at the 0-10m scale (Ω_x_ > 1), and the degree of aggregation was significantly higher in the younger sub-population, consistent with the expectation of increased dispersal limitation over time (Fig 1b).

**FIGURE 1.**
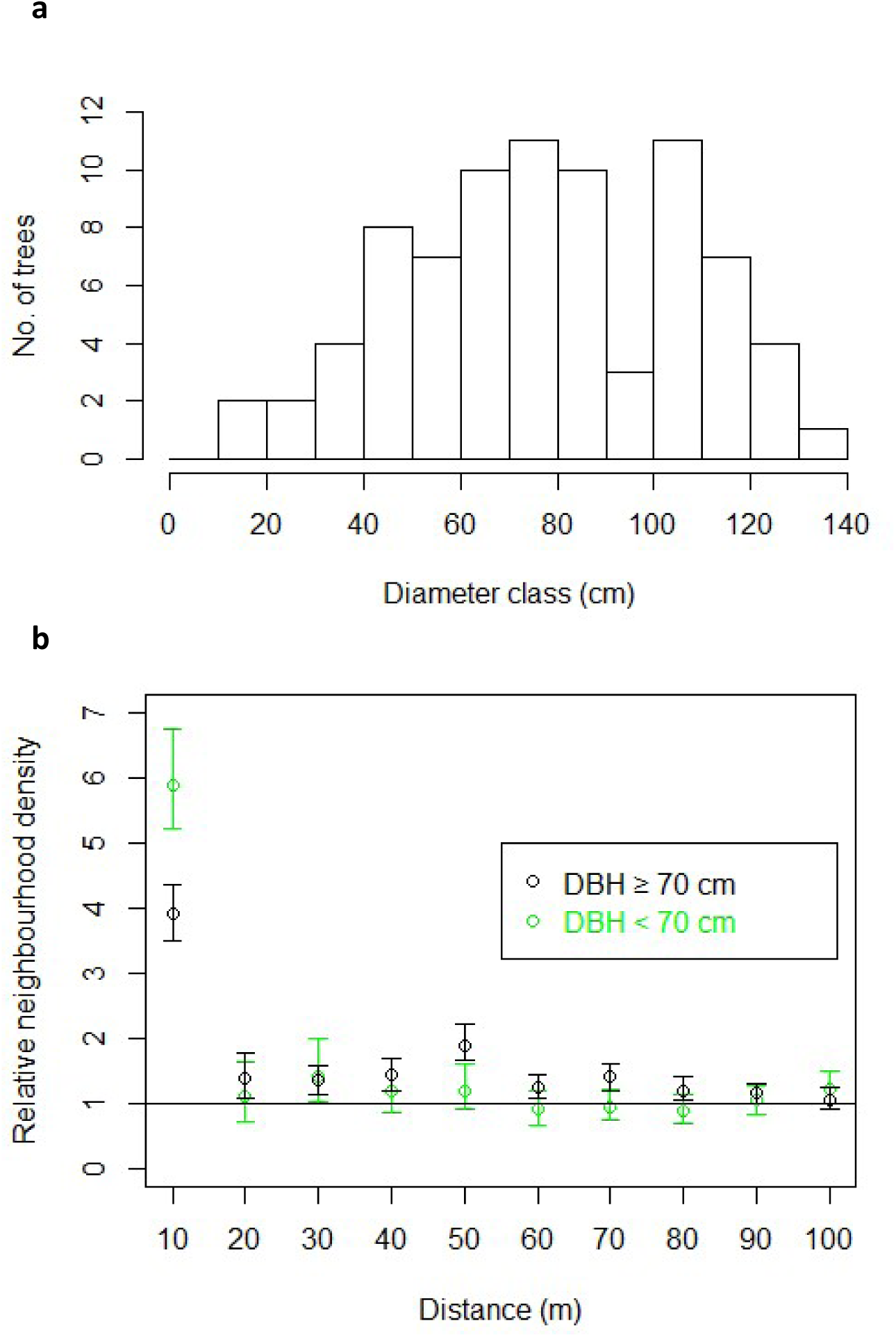
Demographic and spatial distribution of *E. zwageri* adult trees (DBH ≥ 10 cm) in the 4-ha plot. **a)** Population size distribution with a unimodal shape and a mode at the 70 cm class. **b)** Spatial aggregation measured as the mean relative neighborhood density in 10m annuli each focal tree, standardized by mean population density. The population was divided into the older(DBH >70cm) and younger (DBH < 70 cm) sub-populations. Bars represent 95% bootstrapped confidence intervals.

A total of 495 *E. zwageri* seedlings were recorded in 197 of the 540 5 by 5 m subplots surveyed, with an overall mean density of 367 seedlings ha^−1^. Nearly all appeared to be less than one year old and resulting from the fruiting event in January 2017. Among these, 53.3% had fatal herbivory damage, already dead or highly unlikely to survive. The proportion of fatal herbivory damage increased rapidly with conspecific seedling density at subplot level (p < 0.001), predicted to reach 0.97 at 1 seedling m^-2^ (Fig. 2). Fatal herbivory was not significantly associated with vegetation cover (p = 0.09).

**FIGURE 2.**
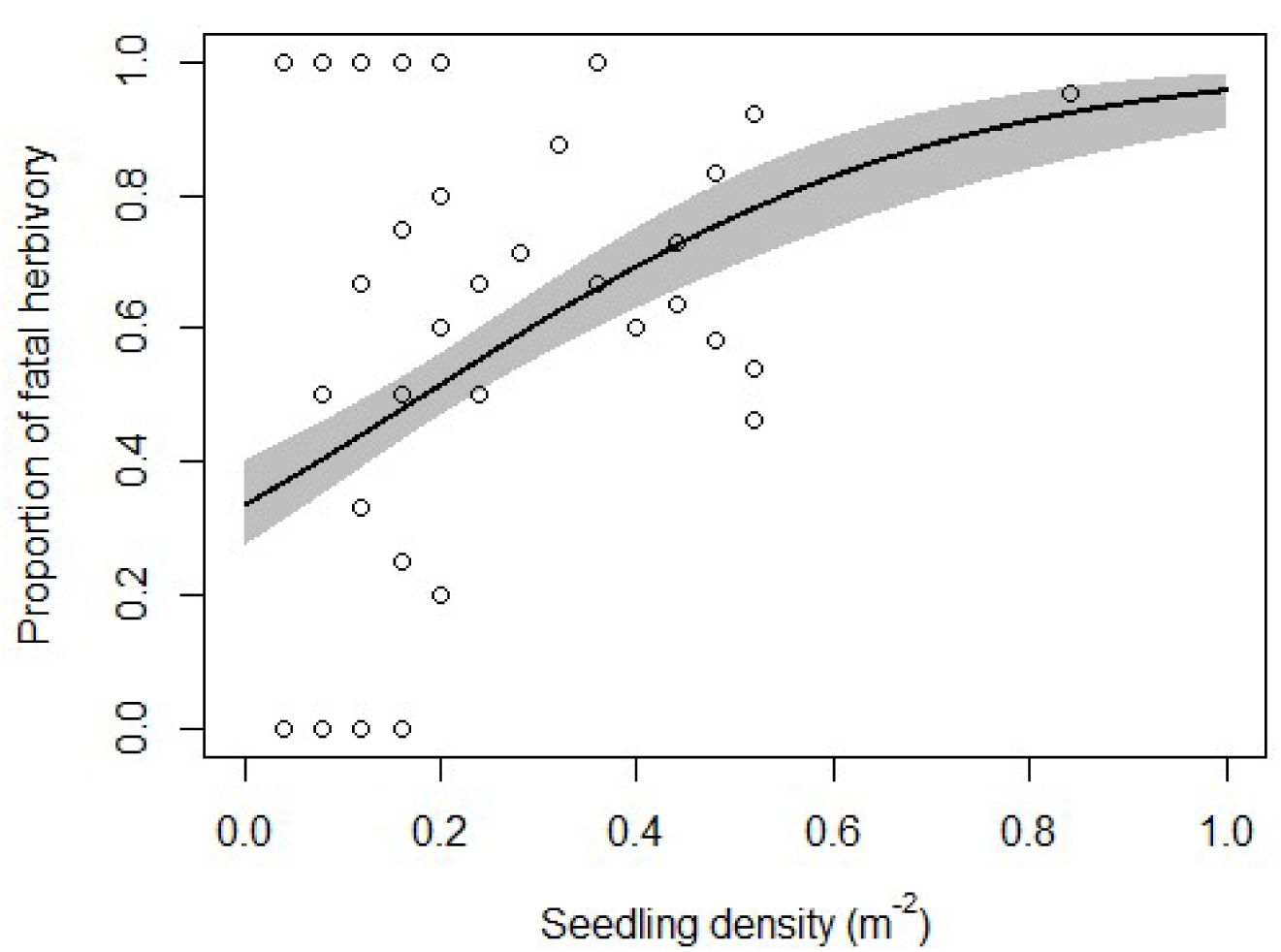
Relationship between the rate of fatal herbivory of *E. zwageri* seedlings and conspecific seedling density at 5×5m subplot level. Line was fitted using a GLM model. Shading areas represent confidence intervals.

## DISCUSSION

The remarkably low abundance of young *E. zwageri* trees indicates that the population of this species is not sustainable in this forest stand (Halpin & Lorimer 2017). *E. zwageri* adult trees were aggregated at the 0-10 m spatial scale, characteristic of tropical trees (Condit *et al.* 2000); a similar aggregation pattern at this scale was observed at genotypical level in a recent study (Shao *et al.* 2017). However, the increased spatial aggregation of *E. zwageri* in younger trees supports the hypothesis of an increasing seed dispersal limitation over time due to the lack of seed dispersers, as shown in another Bornean forest (Harrison *et al.* 2013). Most strikingly, *E. zwageri* seedlings more than one year old (height > 1 m) were almost absent, suggesting complete failure of seedling establishment in recent years. This can be explained largely by the high rate of fatal seedling herbivory, increasing with local aggregation (Fig. 2). These patterns highlight the effect of dispersal limitation and negative density-dependent seedling survival, operating together to influence species spatial distribution (Shao *et al.* 2017).

A survey of 13 other Bornean canopy species found a seedling survival rate of approximately 50% over ten years (Delissio *et al.* 2002). In contrast, we observed an overall survival rate of 46.7% in *E. zwageri* seedlings in less than one year since germination. The actual survival rate may be even lower as we only accounted for dead stems with detectable remains. Using our model as a conservative approximation of the density dependent annual mortality rate and applying it to the surviving *E. zwageri* seedlings at subplot level, we predict that after six years seedling density would drop to merely 3 individuals ha-1, leading to a progressively shrinking demography (Fig. 1a). Seedling height may not reach 2 m in this time (Irawan 2005), and thus they remain susceptible to ground level herbivory.

The heavy herbivory observed here was apparently associated with a single mammalian herbivore, the sambar deer, based on characteristics of the stem damage and reports by local rangers. Adapted to surviving shade conditions for relatively long period, *E. zwageri* seedlings allocate a high proportion of resources towards defense in the form of concentrated tannin (CT) and lignin (Kurokawa *et al.* 2004). This is an evolved mechanism against invertebrate herbivores and pathogens, but mammalian herbivores such as sambar deer have co-evolved to produce a tannin-binding protein in their saliva, and in fact show a preference towards plant species high in CT and lignin (Semiadil *et al.* 1995).

It is possible that at local scale, browsing pressure from sambar deer can influence *E. zwageri* regeneration. A generalist grazer and browser, the sambar deer appears to benefit from intermediate forest disturbance, preferring forest edges and selectively logged forest due to the increased availability of phytomass on the ground (Corlett 2007, Meijaard & Sheil 2007). A recent study in our research area found that mammalian herbivores showed a 19% increase in abundance in logged forest compared to unlogged forest and, in particular, the abundance of sambar deer more than doubled (Wearn et *al.* 2017). There are few large predators in Borneo, and predation pressure on this species here predominantly comes from hunting (Bennett et *al.* 2000). It is likely that in protected areas where hunting is prohibited the sambar deer has enjoyed predator release. Our study site is in a protected area adjacent to a large buffer zone consisting of selectively logged forests. The sambar deer population here therefore have probably benefited from both favorable habitat and the absence of hunting. As an interesting contrast, sambar deer population was low in another Sabah forest (Kabili-Sepilok) (Ross et *al.* 2013), and greater densities of young *E. zwageri* trees were recorded, on average 49 stems with DBH 5 – 10 cm per 4-ha (Qie and Nilus, unpublished data).

Our results shine a spotlight on an iconic tree species E. *zwageri.* We show the intensified interaction between this native tree and a native mammalian herbivore resulting from changes in the megafauna community and forest landscape in Borneo. Anthropogenic forces on the forest ecosystem in recent centuries seem to have eradicated its long-distance seed dispersers and in some places favored its seedling herbivores. Where these impacts co-occur they may lead to an impending regeneration failure of E. *zwageri* at local scale, as we are starting to see in this population.

E. *zwageri* requires active conservation intervention and efforts should focus on its most vulnerable seedling stage. Once survived through this recruitment bottleneck, the tall saplings and trees of *E. zwageri* are known to be exceptionally resilient, including being drought- and fire-tolerant (Delmy 2001, Van Nieuwstadt & Sheil 2005). Ex situ propagation of this species has only been carried out on small scales partly due to inadequate supply of seeds and seedlings (Azani *et al.* 2001, Irawan 2012), yet we show that a great number of *E. zwageri* seedlings in natural forests are being lost to high mortality. Our results suggest that at local density of < 0.2 m^-2^ *E. zwageri* seedlings may have optimal survival. Therefore, moderate harvest of *E. zwageri* seeds and seedlings in natural forests (i.e. wildlings) for ex situ cultivation and re-introduction once they reach sapling stage should be feasible. This approach may simultaneously increase *E. zwageri* regeneration and population gene flow, providing a cost-effective way of assisted forest regeneration.

## ACKNOWLEDGEMENT

This study was supported by the Imperial College London MRes Tropical Forest Ecology field course program. We thank the Sabah Biodiversity Centre for access license and the excellent support of staff members at the Maliau Basin Studies Centre.

